# Microalgal Co-Cultivation meets Media Recycling: A Circular Pathway to Serum and Amino-Acid Reduction in Cultivated Meat

**DOI:** 10.64898/2026.02.04.703452

**Authors:** M. Oey, M.L. Schlieker, U.C. Marx, B. Agustinus, D.M. Vargas Reyes, M. Chandar, B. Hankamer, H. P. Lo

## Abstract

Our increasing global population combined with the UN Sustainable Development Goals of zero hunger and good health require greater protein intake *per capita* and higher protein production. Consequently, sustainable food alternatives such as cultivated meat (CM) are urgently required. However, large-scale CM cell-systems face key challenges, particularly high media costs driven by amino acids and the need for ethically-sourced growth factors. Microalgae offer promising solutions, producing high protein yields with all essential amino acids simply from light, CO_2_, water and nutrients or spent CM media.

Here we present *Chlorella* BDH-1 grown in spent CM media waste as a substitute-source for reduced amino acids and fetal bovine serum in cell culture media, enabling a circular strategy through beneficial mammalian cell-algae co-cultivation. We identified optimal algal growth conditions for maximum protein yield and demonstrated that two recycling rounds using industry-derived spent CM media maximize microalgal biomass yield per unit volume of waste media. We obtained algal lysate, determined thermal processing as the most cost-effective and mammalian cell-beneficial approach, and identified consumed lysate components. Compared to standard media, our lysate increased mammalian cell proliferation over 2-fold in reduced serum and amino acid conditions, replacing costly cell media components. We finally closed the loop by demonstrating a synergistic effect of the algal lysate with our co-cultivation – which co-produces algal biomass. The combination boosted mammalian cell proliferation 1.45-fold, conservatively estimating a media cost reduction by ∼66%. These findings establish parameters to advance the field towards cost-effective sustainable circular cell culture systems with applications in CM production and other biotechnology fields requiring large-scale tissue culture.

Technology Readiness:

## 1 INTRODUCTION

Our global population is projected to increase from ∼8.1 to 10 billion people by 2050. In parallel, the United Nations via its Sustainable Development Goals (SDGs) is focused on delivering *No Poverty, Zero Hunger*, and *Good Health* which will collectively require greater protein intake *per capita*. Combined, these goals would translate into an unsustainable doubling of animal-based food products by 2050^[1]^. Consequently, more environmentally sustainable and ethically acceptable food alternatives are urgently required.

### 1.1 Cultivated meat, as large-scale tissue culture, amplifies media cost challenges

Cultivated meat (CM) has gained significant attention as an alternative food source, due to its ability to deliver sustainable, and cruelty free, high quality food products with minimal or no animal involvement. CM is capable of turning resources into meat 3-times more efficiently than chickens (the most efficient animal system), requiring significantly less land (84-99%) and water resources (43-96%)^[2, 3]^ and releasing less pollutants compared to animal-based meat production^[4-7]^. Furthermore, cultured meat offers significant opportunities to reduce Greenhouse gas (GHG) emissions compared to methane producing lifestock^[4, 7]^. Following this food trend, the sale of cell-cultured meat has now been approved for human consumption in Singapore, Israel, the USA, and more recently Australia/New Zealand, highlighting expanding international market opportunities.

CM production offers important benefits to safety, health, food security, animal welfare and the environment. However, climate benefits of CM production hinge on the scale and energy-type employed for production. Currently, the use of highly refined CM growth media poses environmental concerns due to energy intensive purification processes, resource intensive components and high emission supply chains^[5, 8, 9]^. Therefore, substantial work is being conducted to develop sustainable feed and food-grade culture media. Additionally, CM is economically constrained by the high production costs also faced by traditional laboratory tissue culture, but at a magnified scale. Consequently, market access and expansion require that CM production costs are sustainably as well as renewably driven down to cost competitiveness to generate positive economic and environmental benefits^[5, 7, 8]^. Techno-economic analyses indicate that fed-batch and perfusion facility CM production costs range between ∼ US$37-US$51 Kg^−1^ wet cell mass, significantly exceeding the target price of US$25 Kg^−1^ wet cell mass^[10]^. The key challenges to driving down CM costs therefore include the reduction of the dominating operational costs, in particular cell culture media, which comprises 55-95% of the final product costs ^[2, 10, 11]^.

Fetal bovine serum (FBS), a widely used growth media supplement providing hormones, vitamins, and growth factors^[12]^, is the most expensive component of cell culture media, with prices ranging from US$1,400–1,700 per liter, thus limiting both research and larger-scale applications^[13]^. Rising prices, as well as associated ethical concerns due to the annual use of more than 2 million bovine fetuses to produce 800,000 L of FBS, has increased the urgency to reduce FBS use and identify suitable alternatives^[13, 14]^. To circumvent the use of FBS, many scaled tissue culture applications (including CM) resort to the use of serum-free cultivation. However, currently only ∼260 unique mammalian cell types can be cultured in serum-free media, which still requires the addition of recombinant growth factors or FBS-replacements^[13, 14]^. These are often more expensive, comprising up to 98% of the cost of representative serum-free media^[11, 15]^. The second major cost factor is the provision of amino acids, the necessary building blocks for proteins and cell growth, which significantly contributes to operational costs^[3, 10, 11]^. In the laboratory, amino acids are provided in commercially available growth media such as Dulbecco‚s Modified Eagle Medium (DMEM). However, at scale, cost-effective amino acid supply has the potential to significantly drive down media costs^[10]^. Thus, significant research is being conducted to identify sustainable amino acid sources, with protein hydrolysates shown to effectively promote cell proliferation^[16]^.

### 1.2 Carbon-fixing microalgae are a scalable, low-cost protein source

Microalgae include photosynthetic single-cell, green algae that use sunlight, nutrients, and CO_2_ to produce oxygen and biomass. They are an excellent complement to CM production for oxygen provision as recently demonstrated through co-cultivation^[17]^. Additionally, microalgae production absorbs CO_2_. On a mass basis, dry microalgae biomass typically contains approximately 50% carbon. The relatively undemanding cultivation conditions for microalgae allow for easy, low-cost scale-up. However, one of the most valuable attributes of microalgae is their ability to synthesize all essential amino acids from inorganic nutrients^[18]^. In this way they can ‘up-cycle’ less nutritionally valuable substrates such as CM waste media into valuable protein cell media ingredients for more cost-competitive and sustainable CM production^[19]^. Microalgae have already been identified and widely approved as an attractive sustainable protein source for human consumption. Species with high protein content can contain over 40% protein per biomass dry weight, exceeding soybeans and nuts, and have a comparable amino acid profile to soybean or fishmeal with high levels of complex amino acids (e.g. arginine, lysine, cysteine, glutamine, and glutamate)^[18]^.

### 1.3 Our study

Microalgae extracts have been extensively considered to provide mammalian cell media substitutes, either for FBS or amino acid replacement, but not both^[20-28]^. In this study, we sought to establish parameters for an algae-based circular CM system using the microalgae *Chlorella* BDH-1 ^[17]^. We successfully generated algal lysate from microalgae grown in spent CM media as a substitute for both amino acids and FBS in tissue cell culture, either alone but also in combination with algal-mammalian cell co-cultivation^[17]^. First, we identified the best light quality and temperature for high algal protein yields. We then developed a scalable, low-cost method to process algal lysate, making it suitable to support mammalian cell proliferation in media with reduced FBS and amino acids to a level that is comparable to full serum-containing media. NMR spectroscopy and Mass Spectrometry were used to identify consumed lysate components. We finally applied the obtained knowledge to twice-recycle commercial spent CM media for algal lysate production and demonstrated its ability to substitute for media components in co-cultivation, which improved mammalian cell nutrient consumption, while producing more algal biomass^[17]^. Combining the reduction of costly media components with increased performance, we estimated the media cost savings to be ∼ 66%.

This study provides a significant step towards sustainable, cost-effective, microalgal-supported circular CM economy as it shows the capability of CO_2_ sequestering microalgae to turn spent CM media into novel valuable media components, while reducing FBS and amino acid requirements.

## 2 MATERIAL AND METHODS

### 2.1 Cell lines, culture media, and cell maintenance

#### 2.1.1 Microalgae cells

The microalgae strain *Chlorella* BDH-1 was locally isolated and previously identified as a candidate well suited for co-cultivation with mammalian cells^[17]^. This microalgae cell line is maintained either in liquid TAP medium or on TAP plates^[29]^ containing 1% agar under continuous white LED-light (150 µmol photons m^−2^ s^−1^) and ambient CO_2_.

#### 2.1.2 Mammalian cells

C2C12 myoblasts were obtained from the American type culture collection (ATCC CRL-1772) and routinely grown in Dulbecco‚s Modified Eagle Medium (DMEM, Gibco DMEM 11995065, www.thermofisher.com/au/en/home/technical-resources/media-formulation.9.html) supplemented with 10% fetal bovine serum (FBS - Australian Origin, CellSera), 2 mM L-Glutamine (Gibco), and 100 µg mL^−1^ Ampicillin (Novachem) at 37 °C and 5% CO_2_.

### 2.2 Algae cell density, biomass and protein production in different light and temperature conditions

To identify the best conditions for maximum biomass and protein production per mL culture media, the microalgae were grown in liquid TAP medium and exposed to a variety of continuous light qualities (dark, warm white LED; combined warm/cold white LED at 1:1 ratio, red (630 nm), far red (750 nm), infrared (850 nm) and red combined with 750 nm). The light intensity was maintained at 200 µmol photons m^−2^ s^−1^. Cultures were maintained at ambient CO_2_ concentrations for 21 h at 37 °C, or 48 h at room temperature (20 °C). Cell density (cells mL^−1^) was measured using a Neubauer Hemocytometer. Biomass dry weight was determined by drying down 20 mL of algae culture on pre-dried filter paper. The weight was noted before and after 2 days of drying at 80 °C, and the difference was calculated as biomass dry weight. For protein extraction, the algae were harvested (centrifugation for 10 min at 1000 x *g* at 4 °C), washed in sterile water, and the final algae pellet was resuspended in 25 mL sterile water. The cells were then ruptured at 40,000 psi using a CF1 Continuous Flow Cell Disruptor (Constant Systems). The lysate obtained was then centrifuged (15,000 x *g*, 10 min, 4 °C) to remove cell debris. The supernatant was taken for further processing.

### 2.3 Algae cultivation in cultivated meat spent media

#### 2.3.1 Growth, biomass and protein production

Different batches of spent cultured meat media (CM 1 and 2) were kindly provided by the Australian CM company *Magic Valley Pty Ltd*. Microalgae were cultivated in CM 1 and 2, in comparison to two microalgal media as controls (TAP^[29]^ for mixotrophic growth, PCM^[30]^ for photoautotrophic growth) at continuous white light (150 µmol photons m^−2^ s^−1^) at room temperature and 37 °C. Algae growth was measured based on chlorophyll absorption (680 nm) with a spectrophotometric plate reader (Tecan Infinite M Plex). For biomass assessment, microalgae were grown in spent CM 1 and CM 2 media as well as in TAP and PCM algae media continuously illuminated for 48 h at 37 °C and ambient CO_2_. Cells from 20 mL of culture were washed with sterile water, plated on pre-dried filter paper, and the weight difference was measured after 2 days of drying at 80 °C.

#### 2.3.2 Photoautotrophic microalgal cultivation in twice-recycled spent cultivated meat media

First, CM spent media was diluted 1:1 with sterile water and used for mixotrophic algal growth at ambient CO_2_ and 200 µmol photons m^−2^ s^−1^ at 37 ºC. The resulting half strength recycled CM media was then spiked with 5% photoautotrophic algal media^[29]^ for a second photoautotrophic *C*. BDH-1 cultivation round at continuous light illumination, 32 °C and 1% CO_2_ provision. The algae were then harvested and protein extracted as described in section 2.2.

### 2.4 Protein quantification

For protein quantification, a colorimetric detection assay (Thermo Scientific^TM^ Pierce™ BCA Protein Assay Kit) was used according to the manufacturer’s protocol. In brief, 10 µL of albumin standard dilutions or 10 µL lysate sample were added to a 96-well plate. 200 µL of BCA working reagent was added and the plate incubated in the dark at 37 °C for 30 min. Protein content was quantified by measuring absorption at 562 nm using a plate reader (Tecan Infinite M Plex). Protein concentrations in the respective lysates were determined to be 3.4 mg mL^−1^ for *ALG* lysate (lysate derived from *C*. BDH-1 cultivation in algal media), 2.4 mg mL^−1^ for *CM-Mixo* (lysate derived from mixotrophic algal growth in CM media - 1^st^ CM media recycle round) and 2.2 mg mL^−1^ for *CM-Photo* (lysate derived from photoautotrophic algal growth in CM media – 2^nd^ CM media recycle round).

### 2.5 Processing of microalgal lysate to obtain a solution suitable for cell culture media

To test whether further processing of the algal lysate would improve the replacement of amino acids in standard DMEM growth media, we performed a variety of hydrolysis and protein digestion strategies. After the determination of the protein content, aliquots of *ALG* lysate were either a) processed enzymatically by digestion with 25% or 5% (v/v) Gibco^TM^ Trypsin-EDTA 0.05% (Thermo Fisher) incubated at 37 °C for 2 h (lysate: ES) or 24 h, respectively (lysate: EL); b) hydrolyzed with 0.1 M HCl at 85 °C for 24 h followed by neutralization with 0.1 M NaOH (lysate: AH); c) thermal processing at 85 °C for 24 h (lysate: T). All processed lysates were filter sterilized and stored at −20 °C until use.

### 2.6 C2C12 myoblast experiments

#### 2.6.1 Viability assays

We used a resazurin-based cell viability assay as a measure of mammalian cell proliferation^[31]^. 50 µL of sterile 100 mM resazurin solution was added to 500 µL DMEM media and cells were incubated for 3 h at 37 °C in the dark. Resazurin fluorescence was measured with a plate reader (Tecan Infinite M Plex) at 590 nm using 560 nm excitation.

#### 2.6.2 C2C12 cultivation with microalgal lysate

To assess whether C2C12 myoblasts could utilize algal lysate to proliferate, a basal medium (BM) [1.8 mM CaCl_2_, 2.48 µM Fe(NO_3_)_3_·9H_2_O, 0.81 mM MgSO_4_, 5.3 mM KCI, 44 mM NaHCO_3_, 110 mM NaCl, 0.9 mM NaH_2_PO_4_·H_2_O, 25 mM D-Glucose, 15 mg L^−1^ Phenol Red, 1 mM Sodium Pyruvate, 100 µg mL^−1^ Ampicillin] was prepared. Standard growth medium (GM) consisting of DMEM growth medium supplemented with 10% FBS, 2 mM L-Glutamine, and 100 µg mL^−1^ Ampicillin, was used as control medium. C2C12 myoblasts (10,000 cells per well) were seeded into 24-well plates with 500 µL GM and incubated for 24 h in a standard cell culture incubator (37 °C, 5% CO_2_). The media was then replaced with 500 µL of BM containing 50%, 25% or 12.5% GM (5%, 2.5% and 1.25% FBS, respectively), or 100% GM control media. Cell viability was assessed 48 h after cultivation in each of the dilutions alone or after the supplementation of 100 µL unprocessed lysate (∼250 µg protein) or the supplementation of 100 µL PBS as control to each of the dilutions.

The optimal lysate processing strategy was evaluated by adding 30 µg, 75 µg, 100 µg or 200 µg of differently processed lysates (ES, EL, AH or T) to the BM media containing different percentages of GM. Algal biomass obtained from cultivation in spent CM media was thermally processed, and 200 µg lysate was used for mammalian cell culture experiments, as described above. C2C12 cell viability was measured using a resazurin assay (Fig. 2A).

#### 2.6.3 C2C12 myoblast differentiation after cultivation in algal lysate from spent media

To assess the differentiation of C2C12 myoblasts into myotubes, 40,000 cells were seeded into each well of a 24-well plate and cultured for 3 days in the presence of 200 µg *CM Photo* lysate in different media conditions (100%, 50%, 25%, or 12.5% GM). GM without the addition of any algal lysate was used as a control. Once the cells had reached confluence, the medium was replaced with DMEM supplemented with 2% horse serum to induce differentiation. The cells were then incubated for four days at 37 °C and 5% CO_2_. Images were captured on an AMG EVOSfl microscope using a 4× objective.

#### 2.6.4 Co-cultivation of mammalian cells and algae with lysate

Co-cultivation of C2C12 myoblasts with *C*. BDH-1 was performed according to Oey, et al.^[17]^. To this end, 10,000 C2C12 cells per well were seeded into 24-well plates with 500 µL of 100% GM. The plates were then incubated for 24 hours in a standard cell culture incubator (37 °C with 5% CO_2_) to allow the cells to adhere. The medium was then replaced with 500 µL of 50% GM diluted in BM, to which 50 µg *CM-Mixo* (2.4 mg mL^−1^ protein) or 100 µg *CM-Photo* (2.2 mg mL^−1^ protein) lysate was added, resulting in a final well volume of ∼520 and ∼550 µL, respectively. For co-cultivation, 150 µL *C*. BDH-1 cells in 50% GM was added to each trans-well resulting in a ratio of mammalian cells:algae 1:20^[17]^. For the control experiments without co-cultivation, 150 µL 50% GM was added. The cells were incubated at 37 °C, 5% CO_2_, with continuous illumination of 100 µmol photons m^−2^ s^−1^, and the metabolic activity of the cell population was measured as described in 2.7.

### 2.7 Identification of consumed algal lysate components

To allow for analysis and comparison of components before and after microalgal lysate supplementation, 25% GM samples supplemented with 100 µg microalgal lysate (protein concentration 2.2 µg µL^−1^) were taken immediately before cultivation with C2C12 myoblasts, as well as after 48 h of C2C12 cell cultivation. Medium from cells grown in 25% GM without microalgal lysate were used as a control. To account for sample variation, triplicate samples were pooled and used for NMR and MS analyses.

#### 2.7.1 NMR Spectroscopy

NMR samples were prepared by mixing 160 μL samples with 20 μL 1.5 M potassium phosphate buffer pH 7.4 and 20 μL of 1 mM sodium 2,2-dimethyl-2-silapentane-5-sulfonate (DSS) as a chemical shift reference in D_2_O, yielding a final sample volume of 200 μL with final concentrations of 100 μM DSS, and 10% D_2_O. Samples were transferred into 3-mm NMR tubes for measurement. 1H NMR spectra were recorded using a Bruker 900 MHz Avance NEO NMR spectrometer (Bruker Biospin, Rheinstetten, Germany) operating at a ^1^H frequency of 900.13 MHz and equipped with a 5 mm z-gradient triple resonance cryoprobe (TCI) and a chilled SampleJet sample changer. For each sample, a 1D Carr-Purcell-Meiboom-Gill (CPMG) spectrum was acquired at 298 K with the cpmgpr1d pulse sequence [RD-90°-(τ-180°-τ)n-acq] (Bruker Biospin pulse program library). The transmitter frequency was set to the frequency of the water signal, and water suppression was achieved by continuous wave irradiation during the relaxation delay (RD) of 4.0 s. A fixed spin−spin relaxation delay 2nτ of 80 ms (τ = 500 μs) was used to eliminate the broad signals from high molecular weight analytes. After 4 dummy scans, 128 transients were collected into 65,536 data points using a spectral width of 19,8 ppm, leading to a total experiment time of 13 min per spectrum. All spectra were processed using TOPSPIN version 4.3.0 (Bruker Biospin, Rheinstetten, Germany). The free induction decays (FIDs) were multiplied by an exponential window function with a line broadening factor of 0.3 Hz before Fourier transformation, and manual phase and baseline correction. All media spectra were referenced to the glucose anomeric doublet at δ = 5.233 ppm as this approach resulted in optimal overlay of the various metabolite resonances.

#### 2.7.2 Mass Spectrometry

Prior to the analysis, samples were centrifuged at 17,000 x *g* for 10 minutes at 4 °C and the supernatants transferred into a glass insert. One µL of sample was injected for LC-MS/MS analysis. UPLC-QTOF-MS/MS was acquired on Agilent 6545 QTOF equipped with an Agilent 1290 Infinity II UPLC (Zorbax C_8_ RRHD 1.8 μm 50 x 2.1 mm column, eluting with 0.417 mL/min, 2.50 min gradient elution from 90% H_2_O/MeCN to 100% MeCN with a constant 0.1% formic acid/MeCN modifier). General instrument parameters included gas temperature at 325°C, drying gas at 10 L/min, nebulizer at 20 psi, sheath gas temperature at 400 °C, fragmentation voltage at 180 V and skimmer at 45 V. MS/MS analysis was performed for ions detected in the full scan (100-1700 *m/z)* at an intensity above 1000 counts at 10 scans s^−1^, with an isolation width of 4 ∼*m/z* using a fixed collision energy of 35 eV and a maximum of 3 selected precursors per cycle. UPLC-QTOF-(+)MS/MS data were converted from Agilent MassHunter data files (.d) to an mzML file format using MSConvert software. The resulting data was processed using MS-DIAL (version 5.5). Peak detection was conducted with a minimum peak height of 5000 amplitude, and the MS and MS2 tolerances were set at 0.01 Da and 0.025 Da, respectively. An MS/MS abundance cutoff of 20 amplitude was applied, with a retention time tolerance of 0.1 minutes. MS/MS similarity searches were conducted against the MS-DIAL positive mode MS/MS database for both metabolomic and lipidomic analyses. Annotated results were exported in text format for subsequent analysis. Spectra with low scores or absent MS/MS spectra were still included in the analysis.

### 2.8 Statistical analysis

The C2C12 cultivation data were standardized by scaling each experimental 100% GM control to 1. This enabled comparability across all independent experiments. For each independent experiment, the result was estimated as the mean value of the technical replicates. For each independent experiment the standard deviation was calculated to indicate the degree of dispersion or error. The final overall result was then estimated as the mean across all independent experiments. The overall standard deviation across all independent experiments was computed to indicate the error of the final estimate. The standard error of the mean was obtained for the construction of 95% confidence intervals. For biomass and protein determination, algal growth and co-cultivation, results of each independent sample set were averaged, and their standard deviation calculated. Numbers and values of each independent experiment, as well as the overall results, are provided in Supporting Information Tables.

## 3 RESULTS AND DISCUSSION

### 3.1 Algal growth under red light and elevated temperatures produces highest amount of protein and biomass

As photosynthetic organisms, microalgae respond metabolically to environmental factors^[32]^ including not only to light intensity but also to light quality of different wavelengths^[33, 34]^. Genomic analyses have catalogued dozens of established blue-, red-, and UV-sensitive photoreceptors in marine microalgae, suggesting a complex but still poorly understood light-sensing network regulating lipid synthesis, protein accumulation, and cell division^[35]^. Optimization of light and growth conditions to address both high protein yield and high biomass production is therefore crucial to obtain cost-effective production of microalgal protein lysate to replace cell media components.

For this, *C*. BDH-1 was grown in standard algal medium (TAP^[29]^) using several different light qualities at both RT and 37 °C, and assessed for chlorophyll absorption, cell number, biomass and protein content (Fig. 1, Supplementary Fig. 1). Visual observation of the cultures grown at RT showed a slower growth compared to cultures grown at 37 °C, as previously observed with *C*. BDH-1^[17]^, and thus were grown for 45 h, whereas 37 °C cultures were harvested after 21 h (Supplementary Fig. 1), producing the high biomass yields per mL culture (Fig. 1). In line with other studies^[34, 36, 37]^, our experiments also showed that red light of 630 nm wavelength produced high protein yields per mL culture (Fig. 1). These conditions were therefore chosen as the optimum algal growth parameters for the production of microalgal lysate and implemented for all subsequent applications.

**Fig. 1:**
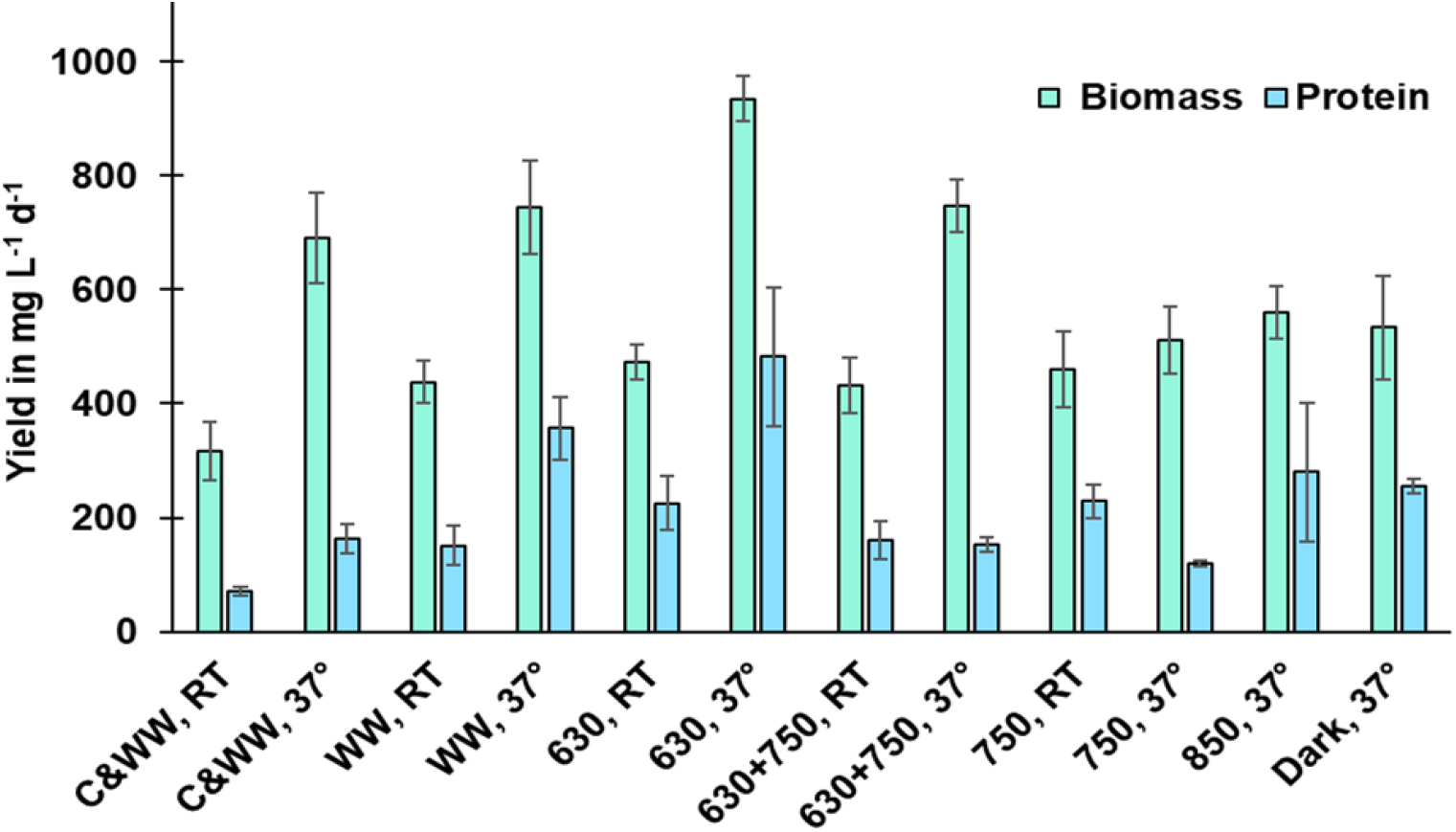
Average biomass dry weight and soluble protein yields (mg L^−1^ d^−1^) obtained from algal cultures grown at different temperatures (RT and 37 °C) and light qualities: cold and warm white (C&WW), warm white (WW), red (630), red and far red combined (630+750), far red (750), near infrared (850) and dark. Mean±SD for biomass was determined from 3 independent experiments; Mean±SD protein content was determined from two independent experiments and a total of 4 sample sets with two replicates each.

### 3.2 Thermally processed algal lysate substitutes for amino acids and FBS in cell media

Several studies have considered using microalgae as a replacement for FBS^[20-24]^, however, these studies relied on the provision of amino acids from standard media.

Using C2C12 cells as a readily available cell-proliferation model, we first determined whether our algae lysate (derived from algae grown in algal medium) could replace both the amino acids and 10% FBS from standard growth media (100% GM). As an indicator, we calculated the concentration of amino acids in DMEM media, with the addition of 2 mM L-glutamine, to be 1.89 mg mL^−1^. For our investigations, Basal Media (BM) with the same composition as standard DMEM growth media but lacking vitamins, amino acids and FBS, was mixed with 100% GM in different proportions to obtain 50, 25 and 12.5% GM (see schematic workflow in Fig. 2A).

**Fig. 2:**
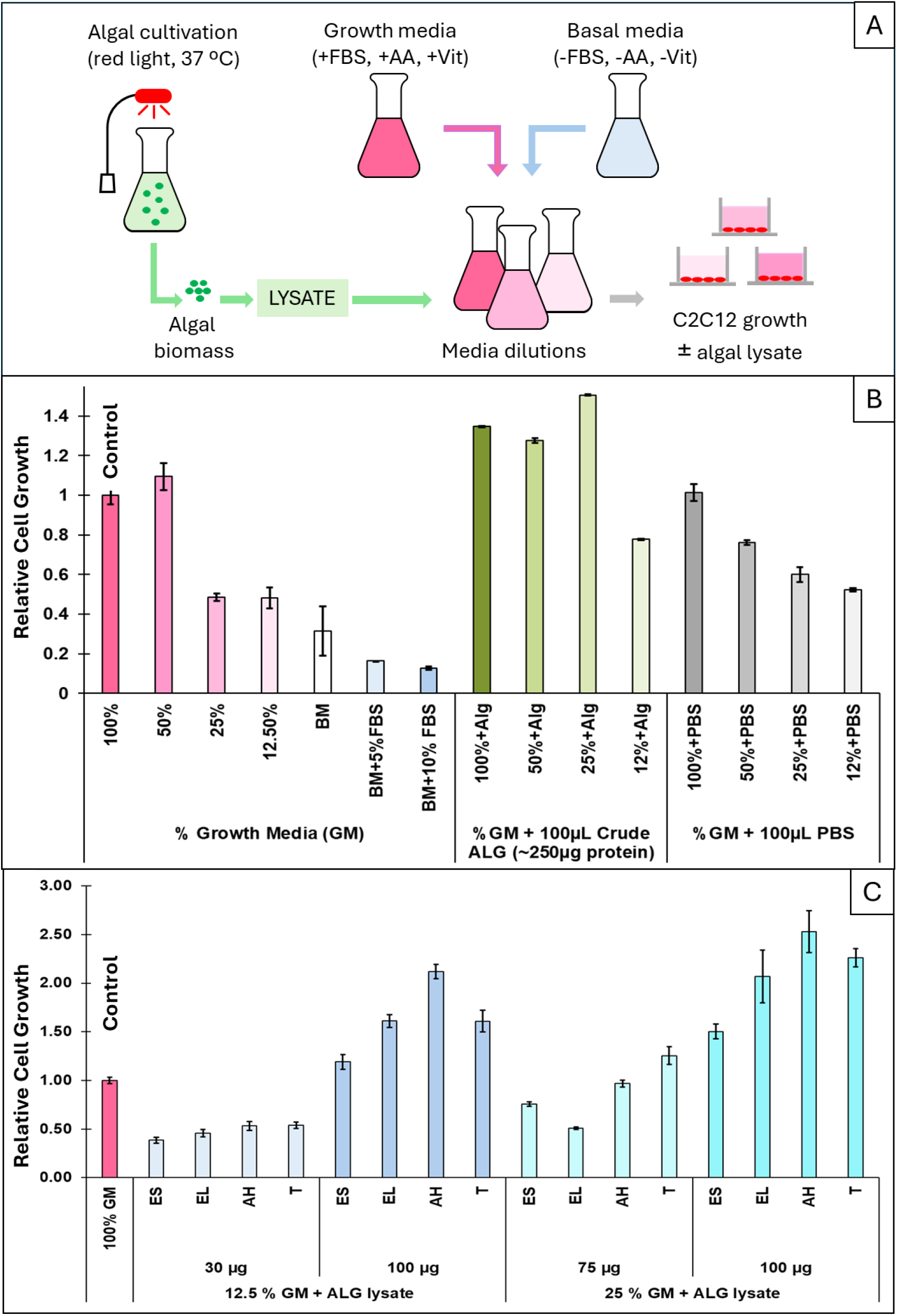
Relative C2C12 cell proliferation in response to addition of microalgal lysate compared to standard conditions (100% GM) after 48 h. **(A)** Schematic illustration of workflow with algal growth in red light, lysate extraction and C2C12 proliferation testing in growth media diluted with basal medium lacking serum (FBS), amino acids (AA) and vitamins (Vit) with or without addition of lysate. **(B)** C2C12 cell proliferation in different percentages of GM (100 [control], 50, 25, 12.5%) with or without the addition of 100 µL of crude, unprocessed algae lysate (*ALG*; ∼250 µg protein) or PBS buffer, as well as pure basal media (BM) with different percentages of FBS (0, 5, and 10%). Mean was obtained from up to 25 independent experiments with up to four technical replicates (Supplementary Table 1). **(C)** C2C12 cell proliferation in standard 100% (control) and 12.5% or 25% of GM with the addition of 30, 75, or 100 µg of *ALG* lysate processed enzymatically (ES: enzymatically short (2 h), EL: enzymatically long (24 h)), acid hydrolyzed (AH) or thermally (T). Mean value was obtained from up to seventeen independent experiments with up to four technical replicates each (Supplementary Table 2). All error bars indicate the 95% confidence interval.

This media was used to cultivate C2C12 cells for 48 h (Supplementary Table 1). As expected, resazurin assays showed a decrease in cell viability down to a minimum of 50% of the control along with the decreasing percentage of GM (Fig. 2B, left). 50% GM itself showed a slightly higher cell proliferation than 100% GM, which could be due to a potentially overall higher starting amount of acidic cell waste products (e.g. lactate) in the 100% GM introduced via FBS as previously described^[17, 38]^. We also prepared BM containing solely 5% or 10% FBS, to assess whether FBS alone could replace the lack of amino acids and determined that neither 5% FBS nor 10% FBS in pure BM could complement the absence of amino acids. Unexpectedly, these conditions performed even less favorably than BM alone, as cells neither proliferated nor maintained viability, showing a reduction of over 80% compared to 100% GM (Fig. 2B left).

We next assessed whether crude, unprocessed algae lysate was able to replace the reduced amino acid and FBS content. *C*. BDH-1 cells grown in TAP medium and red light at 37 ºC were resuspended in water, disrupted, centrifuged and the resulting supernatant was taken as crude algal lysate (termed *ALG* hereafter). 100 µL of crude *ALG* lysate (∼250 µg of unprocessed protein; Fig. 2B, middle) was added to each well (containing different proportions of GM/BM) and C2C12 cell proliferation assessed after 48 h. Interestingly, crude *ALG* enhanced cell proliferation in 100% and 50% GM, achieving a maximum proliferation increase of 35% and 28%, respectively, when compared with standard 100% GM. The biggest difference was detected in cells grown in 25% GM, where cell proliferation was increased by 50% compared to 100% GM. However, in 12.5% GM the crude algae lysate could not restore C2C12 cell proliferation to the same level as achieved in standard 100% GM. We attribute this to cells primarily using single amino acids to synthesize new proteins and sustain proliferation, which might not be possible with crude protein lysate. Lastly, we also tested the addition of 100 µL PBS. As expected, cell proliferation declined when supplemented only with PBS in alignment with decreasing levels of GM (Fig. 2B, right).

We next investigated different *ALG* lysate processing strategies, along with determining the minimum protein content necessary for C2C12 cell proliferation, using 100% GM as the benchmark (Fig. 2C, Supplementary Table 2). Total protein content of lysate was measured, followed by different processing strategies, which were chosen according to ease of handling, availability, and processing cost. We first chose treatment with trypsin, a digestive enzyme that breaks down proteins into smaller peptides in the small intestine, as it is regularly used in cell culture techniques (e.g. for sub-cultivation of adherent cells) and thus readily available. Two different trypsin digest strategies were applied to crude *ALG* lysate: 25% trypsin at 37 °C for 2 h (ES) or 5% trypsin at 37 °C for 24 h (EL). Neither 30 µg or 75 µg of trypsin-digested lysate was able to restore C2C12 cell proliferation to standard levels in 12.5% or 25% GM, respectively. However, 100 µg of trypsin-digested lysate in 12.5% and 25% GM exceeded standard C2C12 proliferation by 60% and 107%, respectively. Overall, the best enzyme-based results were obtained with 100 µg lysate generated with 5% trypsin digest for 24 h (Fig. 2C).

We further processed the crude *ALG* lysate using acid hydrolysis (AH), a conventional strategy in which acidic solutions accompanied by high temperatures are applied to obtain low molecular weight peptides or free amino acids. This strategy was chosen as it is a low-cost procedure applicable on an industrial level. However, in the process, essential amino acids can be destroyed or converted^[39]^. Additionally, the obtained lysates require subsequent neutralization, which causes the formation of salts that can negatively impact the lysate properties. Despite this, acid hydrolysis to process algae lysate for use in cell media has been proposed by Yamanaka, et al.^[26]^ who used 0.25 M, 0.5 M and 1 M HCl in the process. We initially attempted to use 1 M HCl for ALG hydrolysis, however, C2C12 cell proliferation was negatively impacted, potentially due to increased salt concentrations or other factors. Therefore, after considering the requirement of hydrochloric acid with subsequent salt formation from neutralization, we utilized a lower acid content of 0.1 M, considering that these milder conditions would not result in free amino acids. We further tested acid free samples which were only thermally processed (T) at 85 ºC for 24 h alongside the acid containing samples. Neither acid hydrolyzed lysate nor thermally processed lysate showed equal or improved proliferation in 12.5% or 25% GM when applied in low amounts (30 µg). However, supplementation with 75µg of either acid hydrolyzed or thermal lysate in 25% GM showed similar levels of C2C12 cell proliferation compared to standard conditions. 100 µg of both acid hydrolyzed and thermally processed lysates achieved superior C2C12 cell proliferation in both 12.5% and 25% GM (50% - 150% higher cell proliferation compared to standard conditions, Fig. 2C). These results demonstrate that processing plays an important role in making a lysate that can be readily utilized by C2C12 cells. It appears, however, that the provided protein amount was more relevant than the use of acid for lysate processing, as both lysates produced similar proliferation results. This might be due to neither condition being expected to be sufficiently harsh to obtain free amino acids. Thermal processing, which avoids the use of additional acid and neutralization steps or recombinantly produced trypsin, was therefore identified as the best *ALG* lysate processing strategy in terms of cost-effectiveness and ease of handling and thus used for subsequent experiments. This is in line with the recent reporting from Eisenberg et al.^[20]^, who used thermally processed protein extract from *Galdieria sulphuraria*, an extremophilic unicellular species of red algae to replace FBS in cell media. However, although mammalian cell proliferation was performed in full amino acid media, neither of their treatments exceeded cell proliferation in standard growth media. In comparison, the strategy provided here achieved 125% higher (2.25×) cell proliferation using 25% of the standard growth media volume. To produce the same biomass as the standard, the effective media requirement in our system is ∼11% of the standard, corresponding to an up to 90% reduction in media usage and potentially in media cost (not considering separate algae cultivation costs and assuming proportionality).

### 3.3 Identification of consumed media components provided by algal lysate

We next sought to identify the components in the algal lysate that were supporting C2C12 cell proliferation using Nuclear Magnetic Resonance Spectroscopy (NMR) and Mass spectrometry (MS). To do this, we prepared 25% GM media with and without 100 µg thermally processed *ALG* lysate. Samples were taken at 0 h (baseline) and after 48 h incubation with proliferating C2C12 cells. This approach allowed the identification of compounds introduced by the supplemented algae lysate (baseline samples – comparing time points 0 h with or without lysate), as well as those that were consumed by mammalian cells (comparing timepoints 0 h with 48 h).

#### 3.3.1 Nuclear Magnetic Resonance Spectroscopy

We used CPMG NMR spectroscopy to identify compounds that were introduced by the *ALG* lysate. Unexpectedly, low molecular weight compounds detected using CPMG NMR pulse sequences did not show any differences in the observable concentration (above 0.1 mM, the detection limit under the given measurement conditions for a signal arising from a single proton) between samples with or without *ALG* lysate (Fig. 3A and B). Therefore, no specific compounds added by the lysate could be identified using CPMG NMR spectroscopy (Supplementary Fig. 2).

**Fig. 3:**
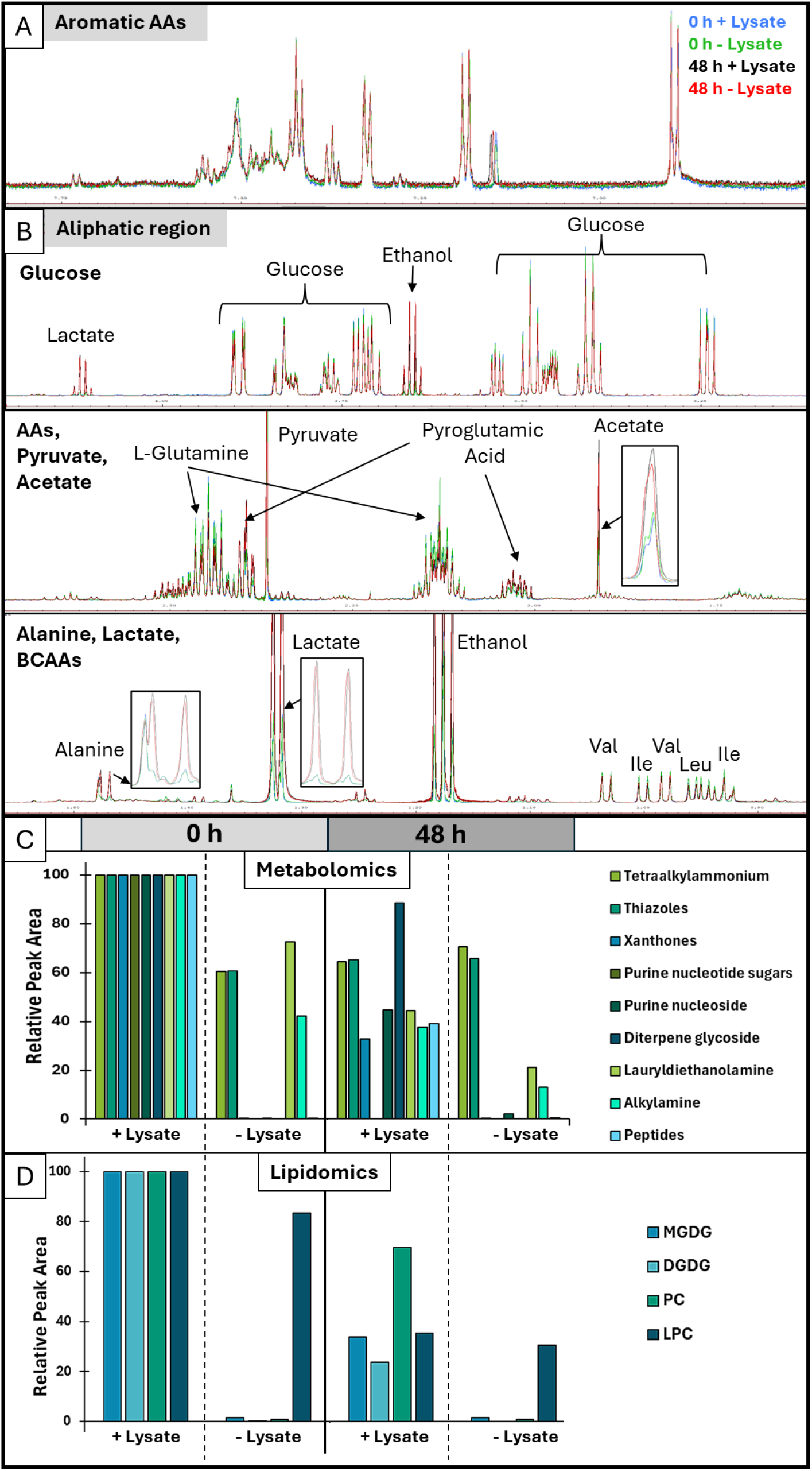
NMR spectroscopy and Mass spectrometry comparison of growth medium with and without 100 µg algal lysate before (0 h) and after 48 h C2C12 cell cultivation. **(A and B)** Overlaid NMR spectra from samples with and without lysate at 0 h and 48 h showing aromatic **(A)** and aliphatic regions **(B)** with peaks identifying amino acids (AAs), glucose, pyruvate, acetate and lactate components. **(C and D)** Mass spectrometry identification of metabolites **(C)** and lipids **(D)** that differ between growth medium with and without algal lysate and that are consumed after 48 h. For comparison, data were normalized to growth medium with lysate at timepoint 0 h (Supplementary Table 3 and 4).

However, differences were observed when comparing the 0 h and 48 h spectra in the aromatic (Fig. 3A) and aliphatic regions (Fig. 3B). As expected, we observed a ∼11% reduction in glucose (from 25 to 22.2 mM) and ∼8% reduction in pyruvate (from 1.8 to 1.65 mM). In contrast lactate increased approximately 9.5-fold (from 2.4 mM to > 50 mM). This implies that the reduction of glucose and pyruvate cannot alone account for the increase of lactate. The sample with *ALG* lysate at higher C2C12 proliferation showed the same glucose and pyruvate consumption, as well as lactate production as the samples without *ALG* lysate, which had lower C2C12 proliferation. This, however, indicated that lysate supplementation supported more C2C12 proliferation per glucose and pyruvate consumption with less lactate production per cell, and vice versa in respective samples without lysate (Fig. 2C).

Under the given measurement conditions, CPMG NMR spectroscopy was able to identify glutamine from the growth media, pyroglutamic acid as a breakdown product of glutamine (for details see Marx and Oey, 2025^[38]^), alanine, as well as the branched chain amino acids valine, isoleucine and leucine. Other amino acids might be present but could not be assigned unambiguously due to chemical shift overlap. As peptide bonds are highly stable under neutral conditions, thermal processing is not expected to release large amounts of free amino acids; this was confirmed by CPMG NMR spectra, which showed that protein or peptides did not breakdown into detectable millimolar levels of single amino acids in the *ALG* lysate. Interestingly, the concentration of branched chain amino acids valine, isoleucine and leucine, essential amino acids that cannot be produced by mammalian cells, was reduced by ∼10% for valine and ∼25% for both isoleucine and leucine after 48 h compared to 0 h samples. The alanine concentration increased approximately 3,65 fold from 0.06 mM to 0.22 mM. We also identified an increase in alanine during the two days of C2C12 cell cultivation, which was previously noted as a suspected by-product of anaerobic glycolysis^[17]^. This indicated a higher alanine secretion per cell from C2C12 cells in medium without algal lysate.

#### 3.3.2 Mass Spectrometry

To complement NMR analysis, untargeted metabolomic profiling using mass spectrometry was conducted to increase sensitivity, and to detect higher molecular weight compounds. As with NMR, samples were taken at 0 h (baseline) and after 48 h incubation with proliferating C2C12 cells, with and without algae lysate. The identified compound classes that were introduced and consumed included choline degradation products, thiazoles (Maillard reaction derivatives), xanthones, purine nucleotide sugars and nucleoside, diterpene glycosides, aminoalcohols, alkyl amine oxides, and peptides (Fig. 3C, Supplementary Table 3). Of these, purine nucleotide sugars and nucleosides provide general substrates for nucleic acid metabolism, whereas plant-based peptides have been reported to promote C2C12 proliferation^[40, 41]^ while generally serving as an amino acid source for cell proliferation. It has further been reported that thiazoles from Maillard reaction are connected to muscle hypertrophy^[42]^.

Additionally, lipid metabolites were assessed, with lipid classes such as monogalactosyldiacylglycerol (MGDG) and digalactosyldiacylglycerol (DGDG), two major structural lipids of photosynthetic membranes in plants and algae^[43, 44]^, phosphatidylcholines (PC), and lysophosphatidylcholines (LPC) identified as being introduced by the algal lysate as well as consumed by the C2C12 cells (Fig. 3D, Supplementary Table 4). While PC and LPC appear to have known effects on muscle cells^[45, 46]^, the effect of MGDG and DGDG is less known.

In combination with the NMR derived data on glucose and pyruvate consumption as well as lactate production, these findings suggested that muscle cells can utilize the energetically dense lipids from the *ALG* lysate as an energy source. It is possible that these feed into the TCA cycle, bypassing glucose and pyruvate consumption, as ATP yields obtained from lipid oxidation can reach 2-3 times those obtained from carbohydrates and amino acids^[47]^. It has further been reported that cardiac muscle cells meet ∼80% of their ATP demands by fatty acids oxidation^[48]^. This is further supported by the observation that growth media without *ALG* lysate provided limited lipid components that were consumed by the C2C12 cells (Fig.3D).

The cause of lactate production, under the conditions tested, remains unclear. Lactate is known to be a product of anaerobic glycolysis and considered as a concentration dependent regulatory signal that switches between glucose and lipid metabolism^[49]^. Therefore, its presence increases the complexity of data analysis beyond this study. Further, the exact connection between lactate and fatty acids remain unclear with research still ongoing despite the phenomenon of fatty acids suppressing glucose catabolism being discovered over 50 years ago^[50]^.

It is also noteworthy that most identified metabolites showed low confidence scores assigned by MS-DIAL based on available MS2 database. For other metabolites, no MS2 spectra was available in the database. This might be due to the thermal treatment producing metabolite by-products that have not been annotated yet and are absent from the current library database.

Thermal processing could also generate more thermostable by-products that are less susceptible to collision-induced dissociation. Therefore, further analysis and method optimization is required for detailed characterization and interpretation, with this study being limited to their identification only (Supplementary Raw data).

### 3.4 *Chlorella* BDH-1 grown in CM waste media produces higher amounts of protein and starch per cell

Cost effective and sustainable scale up is crucial for the CM industry. Spent media should therefore be considered a commodity, not a waste stream. We thus assessed the ability of *C*. BDH-1 to grow in spent media obtained from the Australian CM company *Magic Valley Pty Ltd* for algal biomass and subsequent lysate production. Algal growth was compared to cultivation in standard mixotrophic (TAP) and photoautotrophic (PCM) microalgal media at both RT and 37 ºC. Algal growth in spent CM media was comparable to mixotrophic growth in TAP media at ambient CO_2_, with improved growth at 37 ºC compared to RT (Fig. 4A), consistent with its previously reported temperature preference [17]. Cultures grown at 37 ºC (Fig. 4B) were used for biomass assessment, confirming algal biomass yields obtained from growth in spent CM media (0.61 - 0.76 g L^−1^ d^−1^) to be similar to yields achieved in mixotrophic algal media (0.56 g L^−1^ d^−1^). As expected, photoautotrophic growth produced less biomass (0.11 g L^−1^ d^−1^) (Fig. 4C). Interestingly, the *C*. BDH-1 cell number was reduced in CM media by ∼60% compared to TAP (1.6×10^7^ vs. 4.5×10^7^ cells mL^−1^, respectively). However, the protein content per cell was 3 times higher in spent CM media compared to TAP media (20.1 ng vs. 7.2 ng cell^−1^, respectively) indicating that protein accumulation was increased in spent CM media and prioritized over cell division. Although not the focus of this investigation, we also noted an increase in starch content per cell in the CM media derived samples compared to the TAP cultivated cells (∼13.3 ng vs. ∼2.6 ng cell^−1^, respectively), potentially contributing to a higher biomass weight. This is particularly interesting as the strategy presented here utilizes the soluble fraction for lysate production. However, starch derived from the insoluble fraction could also be processed to serve as a potential source for glucose^[19, 26-28]^.

**Fig. 4:**
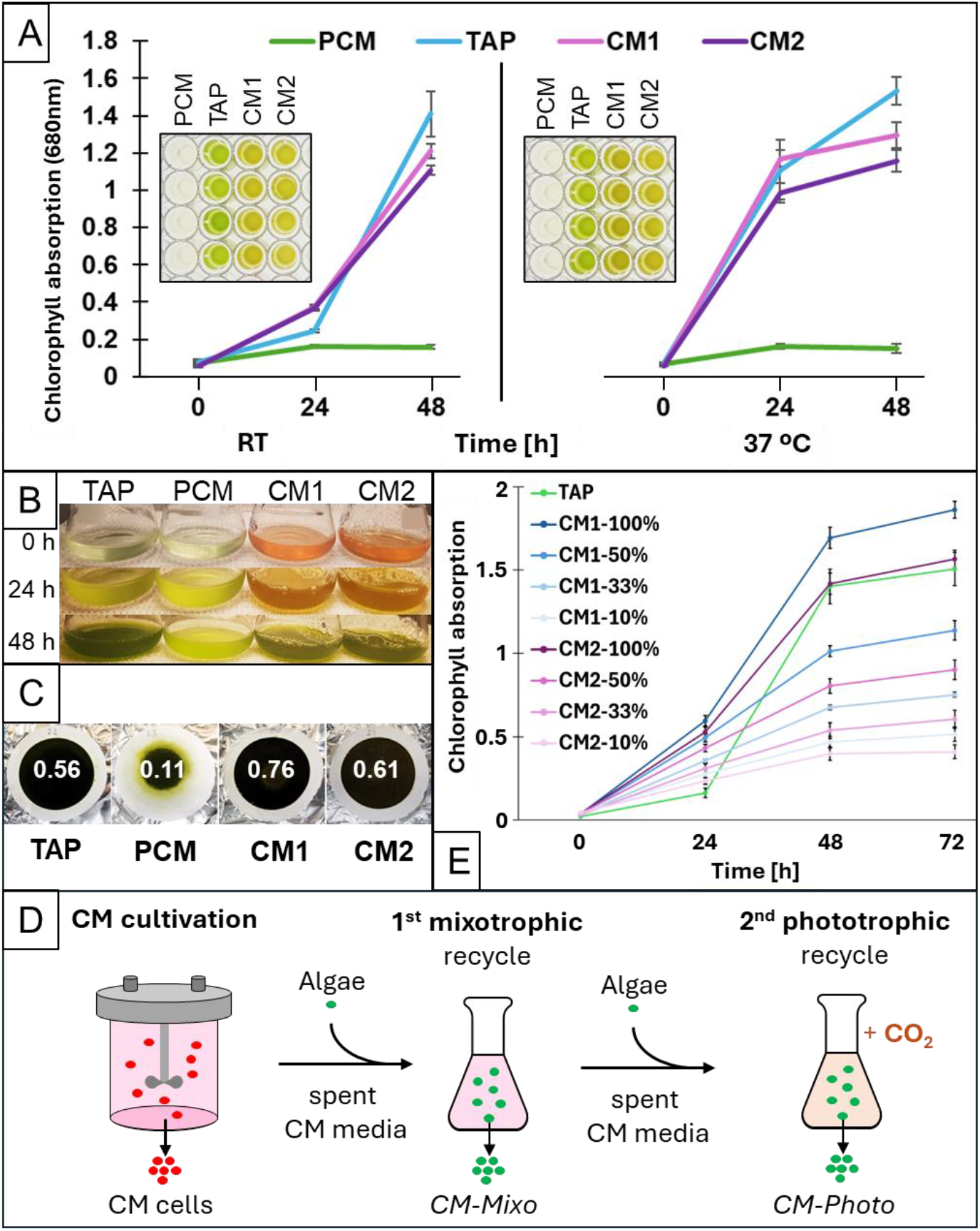
*Chlorella* BDH-1 growth in spent CM media. (**A**) Microalgal growth measured as chlorophyll absorbance at 680 nm in photoautotrophic (PCM) and mixotrophic (TAP) algal media in comparison to spent CM media (1 and 2) at RT and 37ºC. (**B**) Microalgal cultures in TAP, PCM and spent CM medium at 0, 24 h and 48 h. (**C**) Microalgal biomass dry weight in g L^−1^ d^−1^ derived from *C*. BDH-1 cultivated in TAP, PCM and CM spent media. (**D**) Schematic illustration of spent CM media recycling steps for mixotrophic (utilizing organic carbon) and photoautotrophic (utilizing inorganic CO_2_) algal growth and respectively obtained algal biomass for lysates *CM-Mixo* and *CM-Photo*. (**E**) *C*. BDH-1 growth in diluted CM media measured as chlorophyll absorbance (Supplementary Fig. 3). (A and C: Means ± SD, n = 3).

### 3.5 Lysate from twice-recycled spent CM media replaces costly media components

Microalgae can grow mixotrophically by utilizing organic carbon, such as acetate produced as waste from mammalian cells^[17]^, as well as photoautotrophically, using inorganic carbon such as CO_2_. Consequently, microalgal CO_2_ assimilation could provide a useful CO_2_ offset against CM-based emissions, with the potential to lower the overall emissions of a combined microalgae – CM production process. To obtain the maximum amount of microalgal biomass per spent CM media volume for lysate production, we therefore applied two CM spent media recycling rounds – first mixotrophic followed by photoautotrophic algal growth with supplemented CO_2_ (Fig. 4D). The maximum possible dilution of the CM media that would still support optimum algal growth, was established by diluting the spent CM media with sterile water to final concentrations of 50%, 33% and 10% CM media and evaluating *C*. BDH-1 growth (Fig. 4E, Supplementary Fig. 3). Although *C*. BDH-1 chlorophyll content was reduced by ∼30% in 50% spent CM media, a total of 0.8 g L^−1^ d^−1^ biomass was produced, which was comparable to the yield obtained from TAP or pure spent CM medium (Fig. 4C). For the first recycling round, we therefore cultivated the microalgae mixotrophically in 50% diluted spent CM media and processed the extracted lysate thermally, resulting in lysate *CM-Mixo*. Different amounts of *CM-Mixo* lysate were added to C2C12 cells in diluted growth medium and cell proliferation monitored over 2 days (Fig.5A, Supplementary Fig. 4). Supplementation of 100% GM with any tested amount of *CM-Mixo* (25 µg up to 100 µg) lysate increased proliferation by up to 22% relative to 100% GM alone (Supplementary Fig. 4). In 50% GM, as little as 50 µg of *CM-Mixo* lysate was able to establish cell proliferation comparable to standard 100% GM (Fig. 5A). This demonstrates that supplementation with algal lysate from spent CM media can effectively compensate for reduced GM components. We have, however, noted that increasing the amount of *CM-Mixo* lysate beyond 100 µg did not improve cell proliferation and may negatively impact 100% GM (Supplementary Fig. 4).

**Fig. 5:**
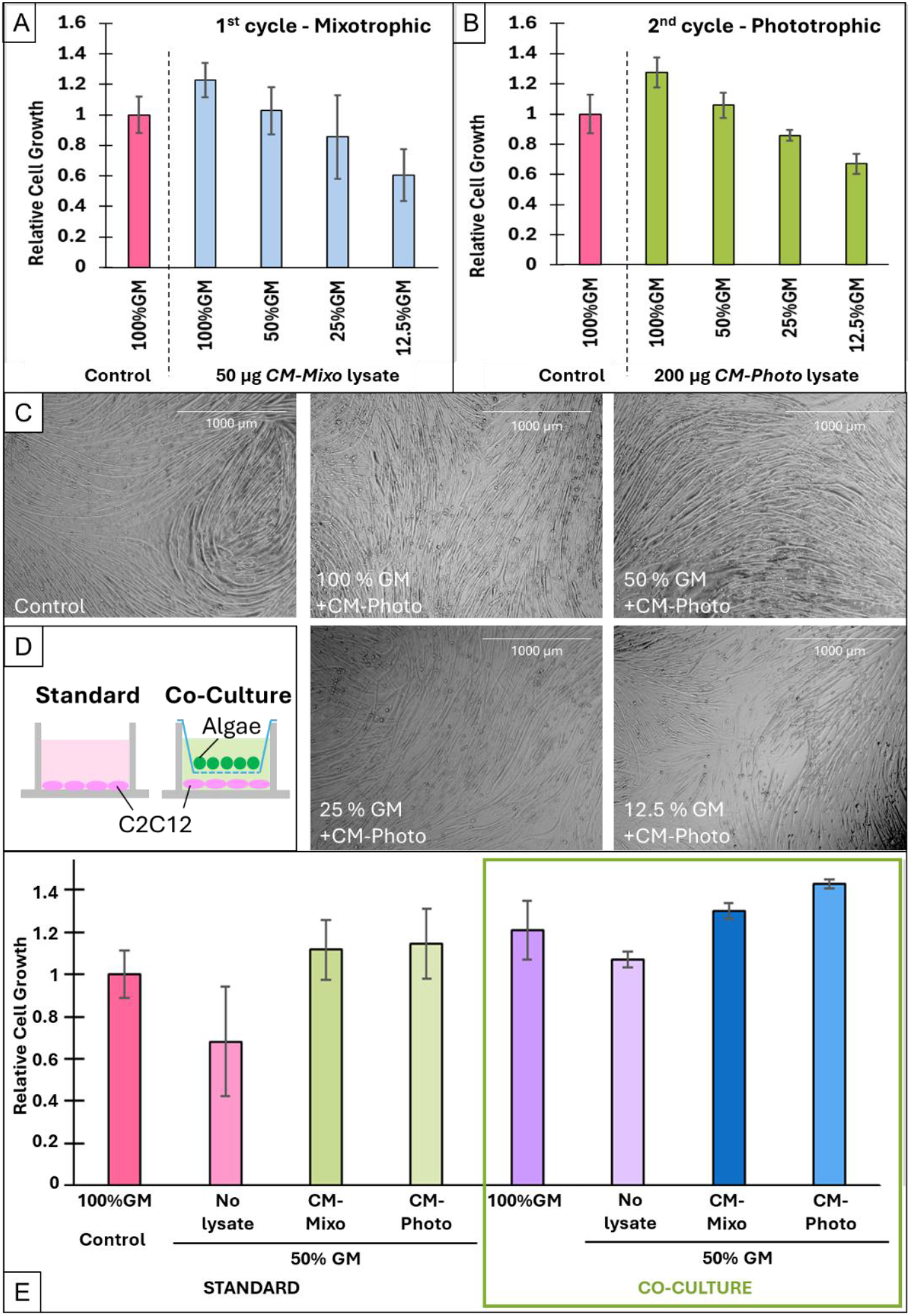
C2C12 proliferation in algal lysate derived from spent CM media. **(A and B)** Relative proliferation of C2C12 cells in 100%, 50% GM, 25% GM and 12.5% GM supplemented with **(A)** 50 µg *CM-Mixo* lysate (1^st^ CM media recycle round, Fig. 4D) or **(B)** 200 µg *CM-Photo* lysate (2^nd^ CM media recycle round; Fig. 4D). Mean values were derived from two independent experiments with up to 6 replicates each (Supplementary Fig. 4 and 6); Error bars indicate the 95% confidence intervals (CI). **(C)** Representative brightfield images of differentiated C2C12 cells after 3 days of cultivation in 100%, 50%, 25% or 12.5% GM with the addition of 200 µg *CM-Photo* lysate followed by differentiation with 2% horse serum for 4 days. Microscope images were taken at 4× magnification. **(D)** Schematic of standard cultivation standard and cell-separation using transwell inserts during co-cultivation (co-culture). **(E)** C2C12 cell proliferation in 100% GM (Control) or 50% GM with or without the addition of 50 µg *CM-Mixo* or 100 µg *CM-Photo* lysate in standard condition (standard) or in co-culture with microalgae *C*. BDH-1 (green box) as described in Oey, et al.^[17]^. Mean±SD, n=3.

For the second recycling round, the CM spent media (after algal removal from the first mixotrophic round) was spiked with 10% photoautotrophic algal medium to replenish inorganic nutrients (Supplementary Fig. 5) and used for photoautotrophic microalgal growth (1% CO_2_, 200 µmol photons m^−2^ s^−1^ illumination, 32 ºC). This twice recycled CM media yielded 0.1 g L^−1^ d^−1^ algal biomass in line with the expected photoautotrophic growth in algal media (Fig. 4C), adding to the overall yield of obtainable microalgal biomass per volume spent CM media. The obtained biomass was thermally processed as previously described to obtain lysate *CM-Photo*. Different amounts of *CM-Photo* lysate were tested with C2C12 cells in diluted GM media to again assess its ability to replace amino acid and FBS in diluted growth media. Like the *CM-Mixo* lysate, the addition of any tested amount of *CM-Photo* lysate to the diluted cell media enhanced the proliferation of C2C12 cells. In 100% GM, supplementation with *CM-Photo* lysate increased proliferation by up to 28% relative to 100% GM alone (Fig. 5B; Supplementary Fig. 6). In 50% and 25% GM, the addition of both 100 µg and 200 µg of *CM-Photo* lysate, supported proliferation comparable to that of 100% GM. This demonstrated that photoautotrophically grown algal lysate derived from spent CM media can effectively substitute for amino acid and FBS components in diluted GM. We further noted that the negative effect previously observed in higher amounts of *CM-Mixo* lysate was absent in *CM-Photo* (Figure 5B, Supplementary Fig. 6). Together, these findings highlight the potential of integrating microalgae cultivation into a medium recycling workflow. Spent media from CM production can support robust microalgal growth through two rounds of media recycling, mixotrophic and photoautotrophic. The resulting lysate can successfully replace amino acids and FBS in mammalian cell media while maintaining high proliferation rates. This strategy enables cell culture resources to be reused and decreases the dependency on costly medium components. Consequently, it enhances the economic and environmental sustainability of CM production.

### 3.6 Differentiation of mammalian cells after proliferation in microalgal lysate medium

In general, cell differentiation is crucial in CM production because it turns generic stem cells into specialized muscle, fat, and connective tissues, giving the final product realistic texture, flavor, juiciness and nutritional value, mimicking real meat‚s complex structure for consumer acceptance^[51]^. To determine whether proliferation with algal lysate under reduced FBS and amino acid conditions would negatively affect C2C12 cells in that regard, we tested their ability to subsequently differentiate. To this end, we cultivated the cells either in 100% GM (Control) or in reduced GM supplemented with *CM-Photo* lysate for 3 days. Subsequently, we induced differentiation and observed the cells for 4 days. In all samples with reduced GM supplemented with algal *CM-Photo* lysate, we observed the formation of multinucleated myotubes (Fig. 5C). This demonstrated that replacement of media components with microalgal lysate supported cell proliferation, while the cells also retained the ability to differentiate.

### 3.7 Closing the loop: algal lysate supports algae-mammalian cell co-cultivation

Finally, we closed the loop towards a circular CM strategy by utilizing the obtained algal lysates to replace GM media components during co-cultivation of microalgae and mammalian cells (Fig. 5D). As previously demonstrated, co-cultivation allows the mammalian cells to more effectively utilize media nutrients, while in parallel algal growth provides oxygen to the system to reduce cellular waste production from the mammalian cells^[17]^. We used lysate obtained from CM spent medium through mixotrophic (*CM-Mixo*) or photoautotrophic (*CM-Photo*) algal growth. 50% GM was chosen as the test medium, as we had previously observed that microalgal co-cultivation alone could only to a limited extent, substitute for reduced FBS (5%) in full amino acid containing growth media^[17]^. We thus aimed to assess whether algal lysate would be able to compensate for this in co-cultivation. Compared to standard 100% GM or 50% GM with lysate only, the combination of both algae-lysate and algal co-cultivation had a synergistic effect leading to higher C2C12 proliferation overall (Fig. 5E). Compared to standard 100% GM without co-cultivation, C2C12 cell proliferation in co-cultivation was 45% higher in samples supplemented with *CM-photo* lysate, and 30% higher in *CM-Mixo* lysate, and accordingly, 25% and 10% higher than in 100% GM with co-cultivation. This demonstrates that algal lysates from spent CM media are able to compensate for the reduction of media components, which have previously hindered cell proliferation in co-cultivation^[17]^. The identified reduction in lipids combined with these results also supports our hypothesis that lipids from algal lysate may act as an additional energy source, as oxygen provided by co-cultivation would enhance lipid oxidation. Our lysate production from algae grown at 37 ºC and in red light is also conveniently complemented by a study from Dani *et al*.^[52, 53]^, who demonstrated that red light had no adverse effect on mammalian cells. Together, these results demonstrate that coupling recycled-media lysate production and use via co-cultivation boosts overall productivity and establishes an efficient, closed-loop cultured meat production workflow. This strategy substantially reduces media costs while enhancing cell performance, highlighting its strong potential for scalable, cost-effective cultured meat bioprocesses.

## 4 CONCLUSION

To our knowledge, this is the first study to demonstrate that microalgal lysate derived from recycled CM media can substitute for media with reduced FBS and amino acids, while concurrently boosting beneficial mammalian cell proliferation through algal co-cultivation. In fact, this strategy obtained 45% higher mammalian cell proliferation than standard growth conditions. Our approach combines microalgal cultivation in CM waste for lysate production - replacing costly amino acids and serum in cell media - with microalgal co-cultivation - for oxygenation, improved nutrient usage by mammalian cells and production of more microalgal biomass - which together improve cell proliferation.

Microalgae extracts have been extensively considered as cell media substitutes, and several studies have evaluated their use as replacement of FBS or amino acids. However, these studies either relied on the supply of amino acids from standard media^[20-25]^, replaced amino acids with algal lysate alongside full 10% FBS provision^[26]^ or demonstrated improved performance only in comparison to control cells grown in deficient media, which likely impaired cell performance^[27, 28]^. Haraguchi and Yamanaka, *et al*. provided insights into the use of algal lysate in conjunction with growth factor secreting cells ^[19, 26]^.

In this study we have a) identified the best *C*. BDH-1 cultivation conditions for high biomass and protein production, b) established cost-effective, large-scale suitable processing strategies to obtain algal lysate that can be c) substituted for amino acids and serum, while producing comparable or superior cell proliferation to standard medium. We have applied NMR spectroscopy and Mass spectrometry to identify compounds provided by the algal lysate that are consumed by the C2C12 cells. Based on our results, we hypothesize that algal lipids are an additional energy source for the C2C12 cells, providing a basis for future investigations and supplementation studies. We also used our algae to first mixotrophically then photoautotrophically recycle spent CM media obtained from *Magic Valley Pty Ltd* and demonstrated that the obtained algal lysate can be used in mammalian cell-algal co-cultivation, to increase mammalian cell proliferation with the concomitant ability to produce new algal biomass in the same reactor.

This demonstrates the feasibility of two circular cost-saving scenarios (Fig. 6) including a two-step approach - separating CM from algal cultivation, as well as a co-culture approach that combines recycled-media lysates with productivity boosting co-cultivation. Assuming a conservative 50% cost savings of costly media components using 50% GM for the latter scenario, in conjunction with 45% increased productivity and no additional algal cultivation costs, we estimate potential media cost reductions up to ∼66%, with waste management savings also not considered. Additionally, our integrated microalgal-CM strategy has the potential to reduce CO_2_ emissions and positively influence the climate dynamics of the overall CM process (Fig. 6). However, further lifecycle analyses are required for a more detailed assessment.

**Fig. 6:**
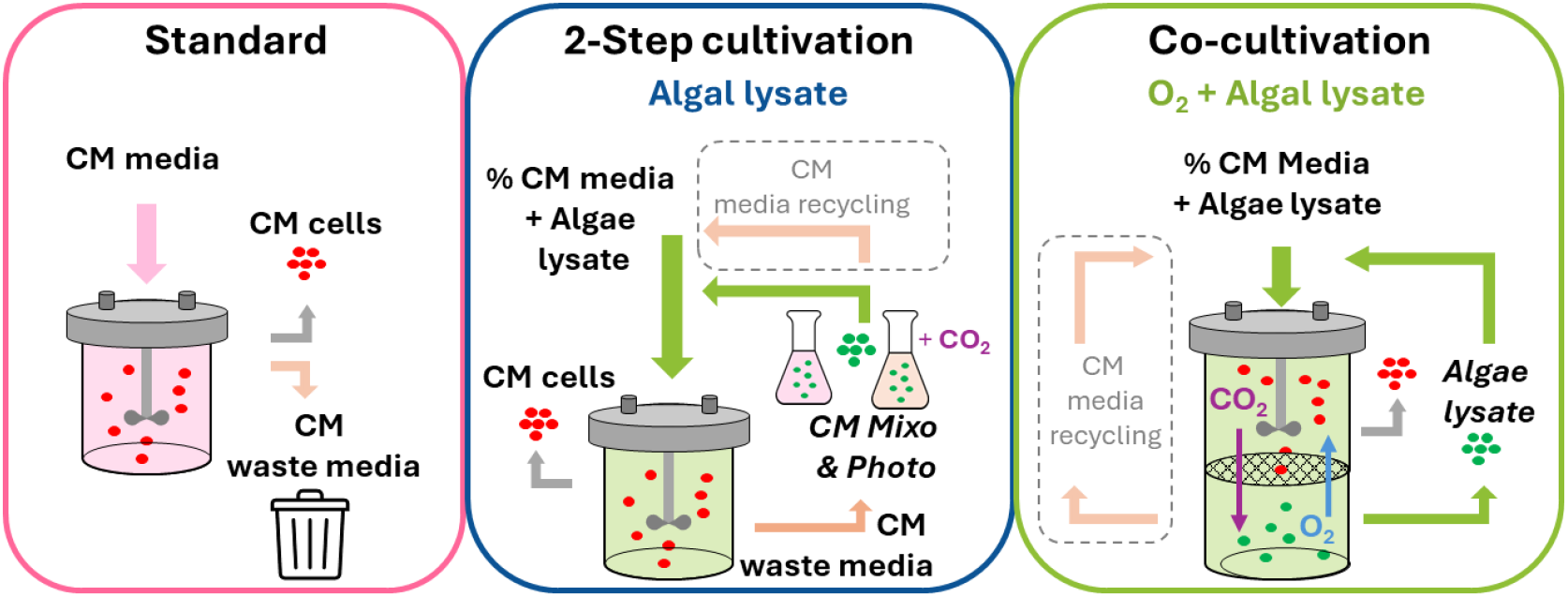
Schematic illustration of standard procedure versus two proposed cost-reducing scenarios including a 2-step cultivation, which separates CM cell proliferation and algal growth, as well as a co-cultivation scenario, in which oxygen producing algae are co-cultivated to improve nutrient usage while producing algal biomass for lysate production. We also included potential media reusage (grey box) as proposed by Haraguchi, et al.^[19]^ to complement our strategy.

Currently, the absence of a suitable scale-up system for co-cultivation (currently limited to trans-well plates) inherently restrains the ability to produce sufficient microalgal biomass from co-cultivation and is thus a focus of future work. We further acknowledge that C2C12 cells, an immortalized cell line used as model in laboratory settings, may not be considered ideal representatives of cells used in commercial CM production and thus are a limitation to our study. However, with the broad range of often IP protected proprietary cell lines being developed in the CM industry, potentially requiring customized optimization in conjunction with their respective proprietary spent CM media, we consider the work presented here a feasibility study providing a path towards cost-effective scale-up of numerous processes.

We show that our strategy enables cell culture resources to be reused, thus decreasing dependency on costly medium components while enhancing production output. Consequently, it has the potential to enhance the economic and environmental sustainability of CM production and other mammalian cell cultivation at scale.

## Acknowledgments

We acknowledge Magic Valley Pty Ltd for the kind provision of their cell culture waste for the performed cell proliferation experiments. We thank Dr Rhia Stone, Dr Robert Chapman and Ashlee Rodd for their help in sample preparation, Dr Gregory Pierens and Dr Thomas Durek for support with NMR sample preparation and measurements as well as Dr Angela Salim and Paayal Kumar for assistance with Mass Spectrometry measurements. We would like to thank Prof. Robert Parton for insightful discussions and constructive comments on the manuscript.

## 6 Supplementary Data

**Supplementary Fig. 1:**
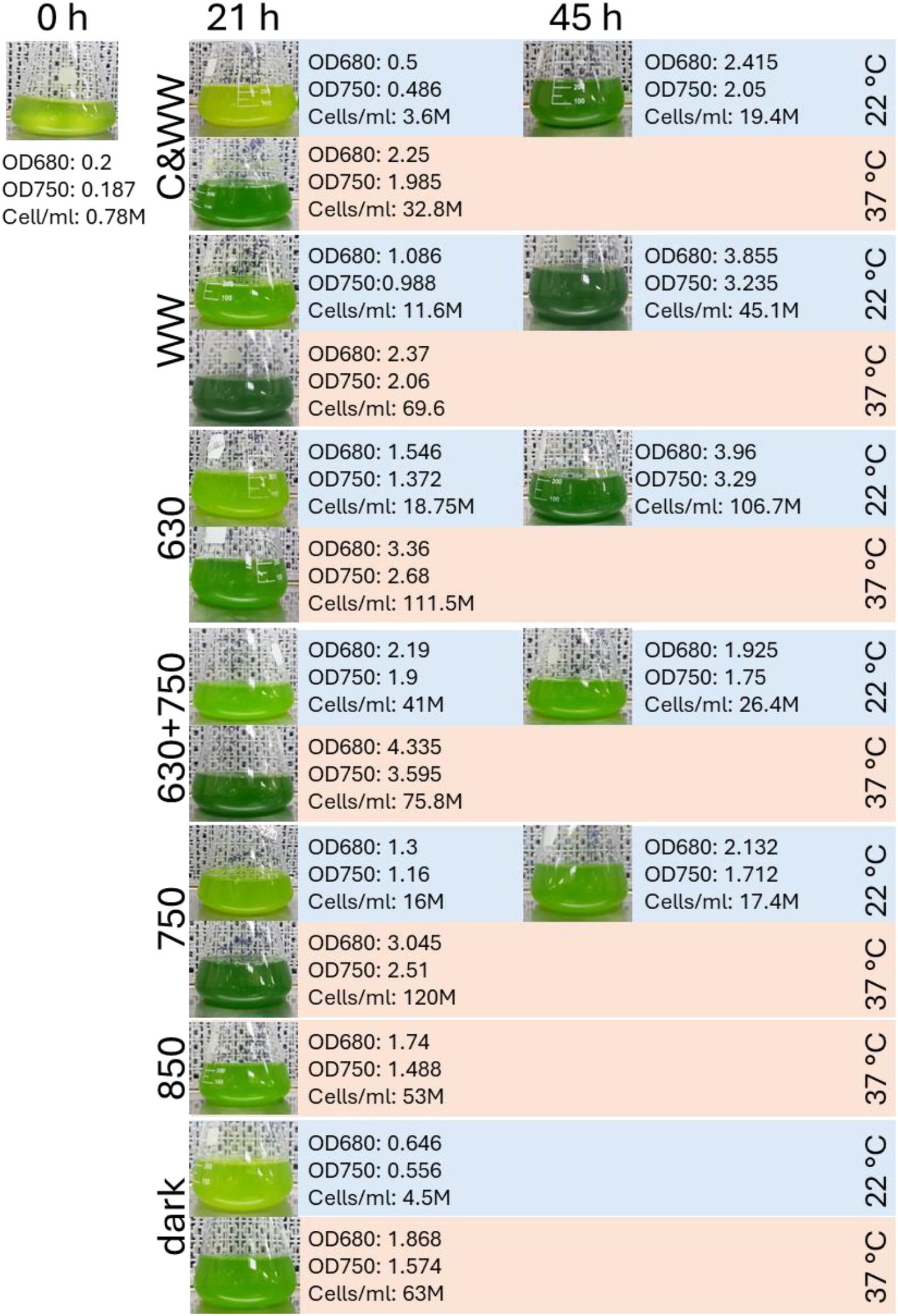
Visual assessment of algae cultures grown in different light and temperatures, along with chlorophyll absorption at 680nm (OD_680_) and cell count per mL.

**Supplementary Table 1:** Relative fluorescent units derived from resazurin assay representing C2C12 proliferation in different percentages of growth medium with or without 100 µL crude algal lysate or PBS.

| RATIOS | % Growth Media (GM) | | | | | | | %GM + 100μL Crude ALG (~250μg protein) | | | | %GM + 100μL PBS | | | |
| --- | --- | --- | --- | --- | --- | --- | --- | --- | --- | --- | --- | --- | --- | --- | --- |
|  | 100% | 50% | 25% | 12.50% | 0% | BM+5%<br>FBS | BM+10%<br>FBS | 100%+Alg | 50%+Alg | 25%+Alg | 12%+Alg | 100%<br>+PBS | 50%+<br>PBS | 25%+PB<br>S | 12%+PBS |
| Data set |  |  |  |  |  |  |  |  |  |  |  |  |  |  |  |
| 1 | 1 | 1.383249 |  |  | 0.378173 |  |  | 1.7081218 | 1.284264 |  |  |  |  |  |  |
| 2 | 1 | 1.823564 | 1.34823 |  | 0.250725 | 0.1828 | 0.09576 | 1.0510737 | 1.4143935 | 2.037725 |  |  |  |  |  |
| 3 | 1 | 0.642937 |  |  |  |  |  |  |  |  |  |  |  |  |  |
| 4 | 1 | 0.87415 |  |  |  |  |  |  |  |  |  |  |  |  |  |
| 5 | 1 | 0.750376 | 0.595489 | 0.52030075 |  |  |  | 1.281203 | 1.1293233 | 0.974436 | 0.7774436 | 1.014 | 0.76 | 0.6 | 0.523308 |
| 6 | 1 |  | 0.946088 | 0.38157753 |  |  |  |  |  |  |  |  |  |  |  |
| 7 | 1 |  |  | 0.86175507 |  |  |  |  |  |  |  |  |  |  |  |
| 8 | 1 |  |  |  |  | 0.1437 | 0.15878 |  |  |  |  |  |  |  |  |
| 9 | 1 |  |  | 0.0998935 |  |  |  |  |  |  |  |  |  |  |  |
| 10 | 1 |  |  | 0.49120971 |  |  |  |  |  |  |  |  |  |  |  |
| 11 | 1 |  |  | 0.43346096 |  |  |  |  |  |  |  |  |  |  |  |
| 12 | 1 |  |  | 0.55351682 |  |  |  |  |  |  |  |  |  |  |  |
| 13 | 1 |  |  | 0.51220126 |  |  |  |  |  |  |  |  |  |  |  |
| 14 | 1 |  | 0.183771 |  |  |  |  |  |  |  |  |  |  |  |  |
| 15 | 1 |  | 0.254184 |  |  |  |  |  |  |  |  |  |  |  |  |
| 16 | 1 |  | 0.214248 |  |  |  |  |  |  |  |  |  |  |  |  |
| 17 | 1 |  | 0.16876 |  |  |  |  |  |  |  |  |  |  |  |  |
| 18 | 1 |  | 0.442391 |  |  |  |  |  |  |  |  |  |  |  |  |
| 19 | 1 |  | 0.438782 |  |  |  |  |  |  |  |  |  |  |  |  |
| 20 | 1 |  | 0.451333 |  |  |  |  |  |  |  |  |  |  |  |  |
| 21 | 1 |  | 0.453449 |  |  |  |  |  |  |  |  |  |  |  |  |
| 22 | 1 |  | 0.442391 |  |  |  |  |  |  |  |  |  |  |  |  |
| 23 | 1 |  | 0.438782 |  |  |  |  |  |  |  |  |  |  |  |  |
| 24 | 1 |  | 0.451333 |  |  |  |  |  |  |  |  |  |  |  |  |
| 25 | 1 |  | 0.453449 |  |  |  |  |  |  |  |  |  |  |  |  |
| Overall Average | 1 | 1.094855 | 0.485512 | 0.48173945 | 0.314449 | 0.1633 | 0.12727 | 1.3467995 | 1.2759936 | 1.50608 | 0.7774436 | 1.014 | 0.76 | 0.6 | 0.523308 |
| Overall Std Dev | 0.117445 | 0.077756 | 0.034699 | 0.07655809 | 0.090119 | 0.0009 | 0.00499 | 0.0030075 | 0.0105263 | 0.003008 | 0.0021266 | 0.021 | 0.01 | 0.01914 | 0.004253 |
| Overall SEM | 0.023489 | 0.034773 | 0.008959 | 0.02706737 | 0.063724 | 0.0006 | 0.00353 | 0.0017364 | 0.0060774 | 0.002127 | 0.0021266 | 0.021 | 0.01 | 0.0191 | 0.00425 |
| 95% CI | 0.046038 | 0.068156 | 0.01756 | 0.05305205 | 0.124898 | 0.0012 | 0.00691 | 0.0034033 | 0.0119116 | 0.004168 | 0.0041682 | 0.042 | 0.01 | 0.03751 | 0.008336 |

**Supplementary table 2:**
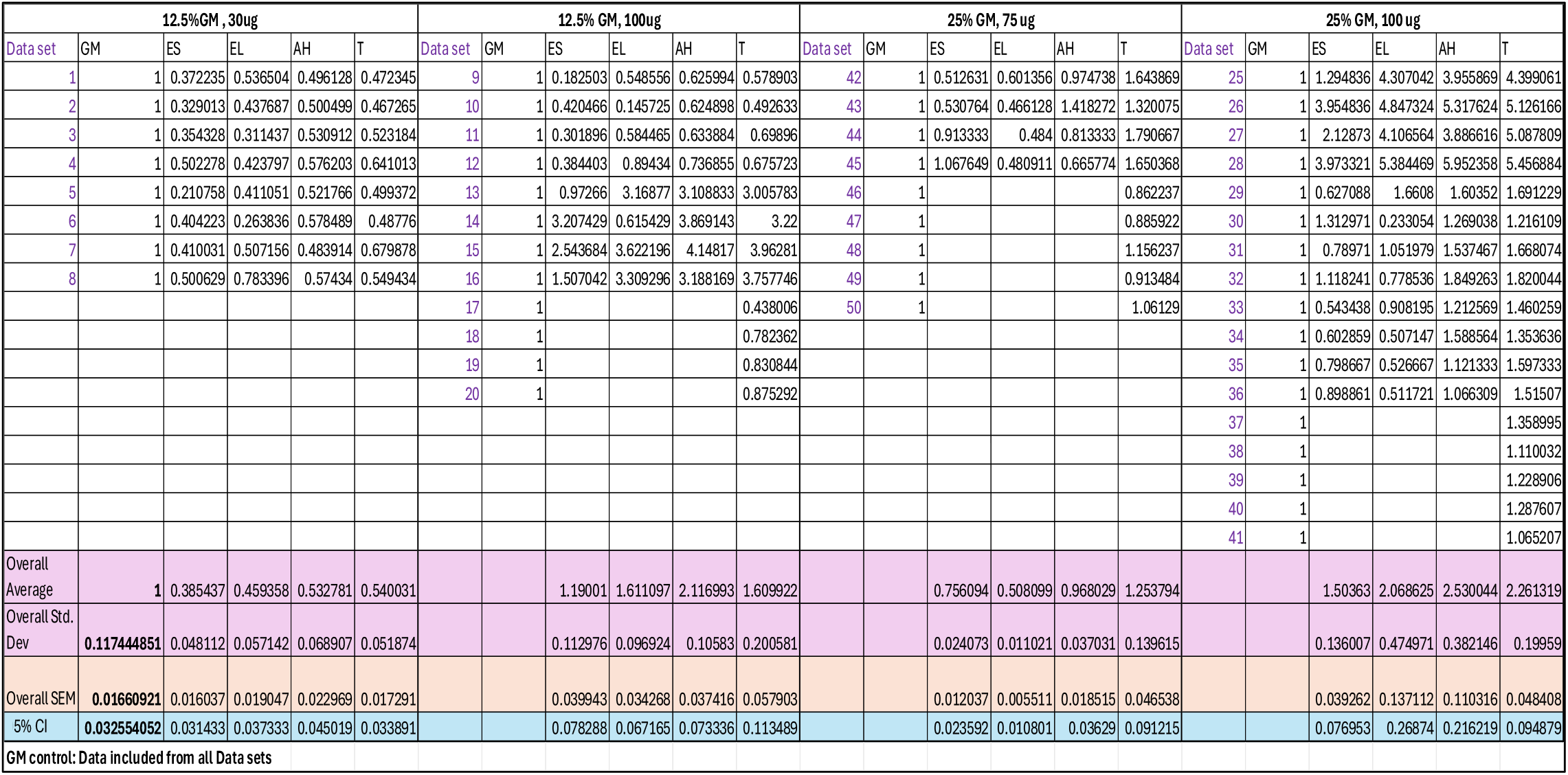
Relative fluorescent units derived from resazurin assay representing C2C12 cell proliferation in different percentages of growth medium with various amounts of differently processed algal lysates (enzymatic short (ES= 2 h), enzymatic long (EL = 24 h), acid hydrolysis (AH) and thermally processed (T).

**Supplementary Fig. 2:**
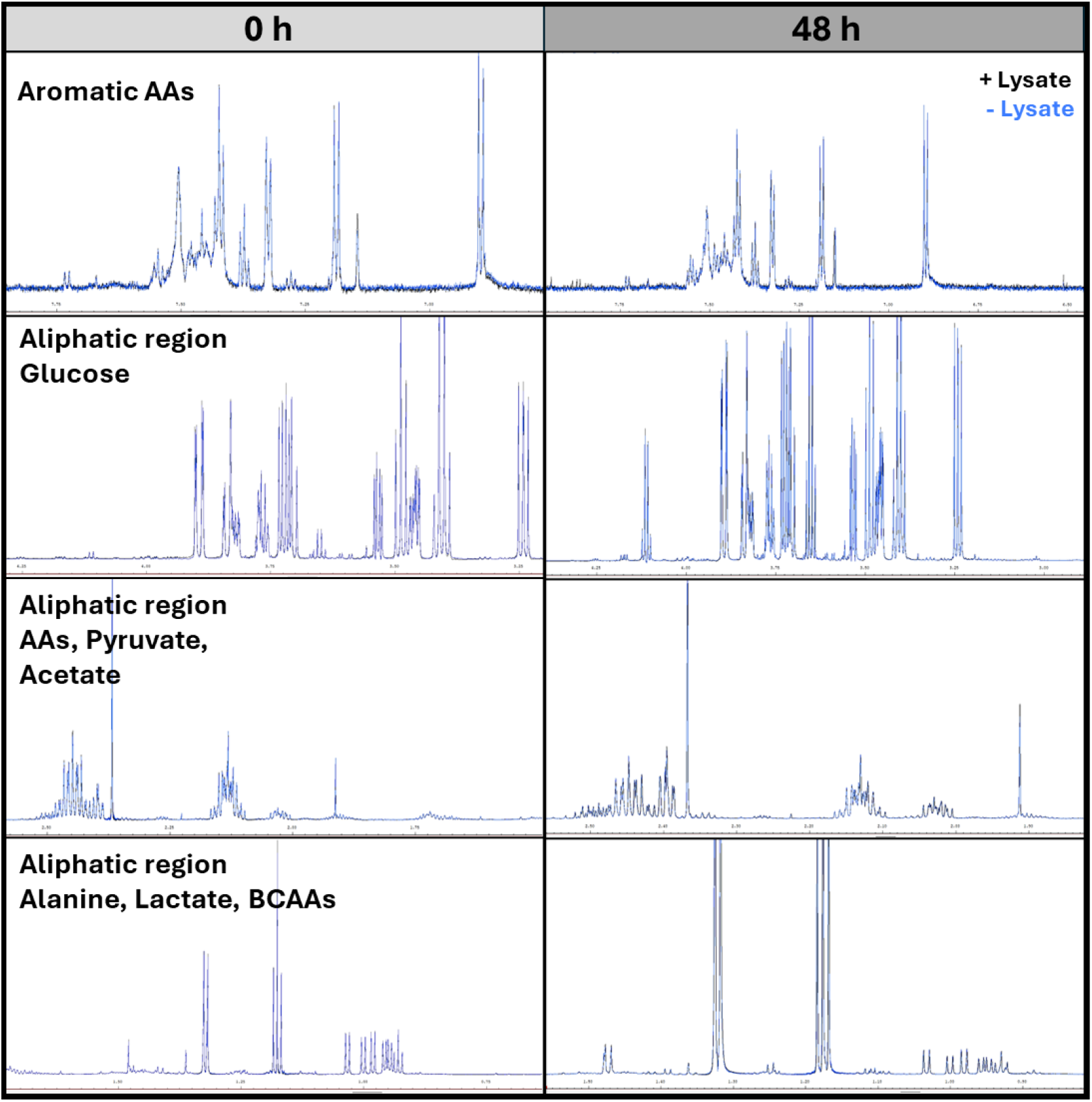
NMR spectra from samples with and without lysate at 0 h and 48 h showing aromatic and aliphatic regions.

**Supplementary Table 3:**
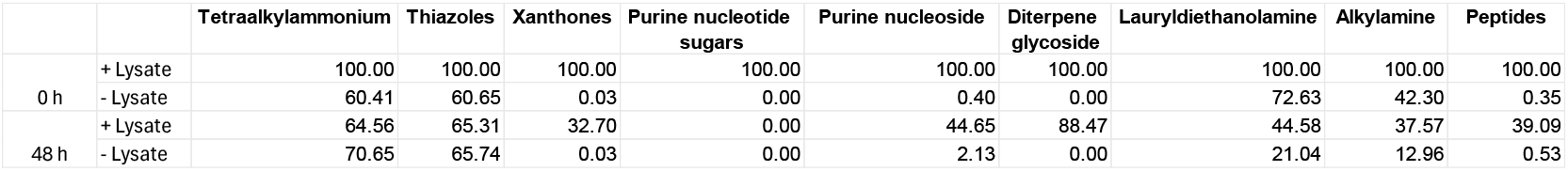
Metabolomics data of the compounds identified to be provided by the lysate as well as being consumed by the C2C12 cells. Values represent the percentage (%) of compound area relative to samples supplemented with *ALG* lysate (∼ 100 µg of algal protein) at t = 0 h.

**Supplementary Table 4:** Lipidomics data of the compounds identified to be provided by the lysate as well as being consumed by the C2C12 cells. Values represent the percentage (%) of compound area relative to samples supplemented with *ALG* lysate (∼ 100 µg of algal protein) at t = 0 h.

|  |  | MGDG | DGDG | PC | LPC |
| --- | --- | --- | --- | --- | --- |
| 0 h | + Lysate | 100 | 100 | 100 | 100 |
|  | - Lysate | 1.37 | 0.24 | 0.85 | 83.45 |
| 48 h | + Lysate | 33.79 | 23.70 | 69.57 | 35.22 |
|  | - Lysate | 1.46 | 0.00 | 0.79 | 30.60 |

**Supplementary Fig. 3:**
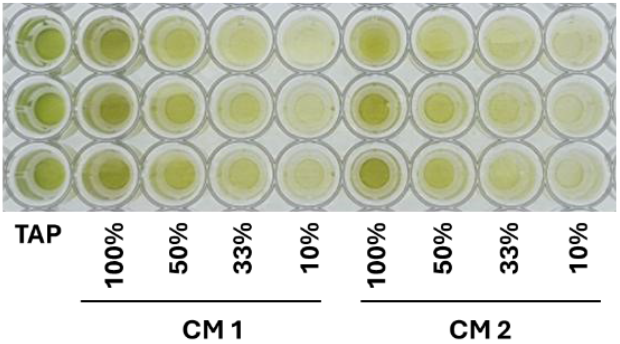
Visual example of *C*. BDH-1 growth in diluted spent CM media at 72 h complementary to Fig. 4

**Supplementary Fig. 4:**
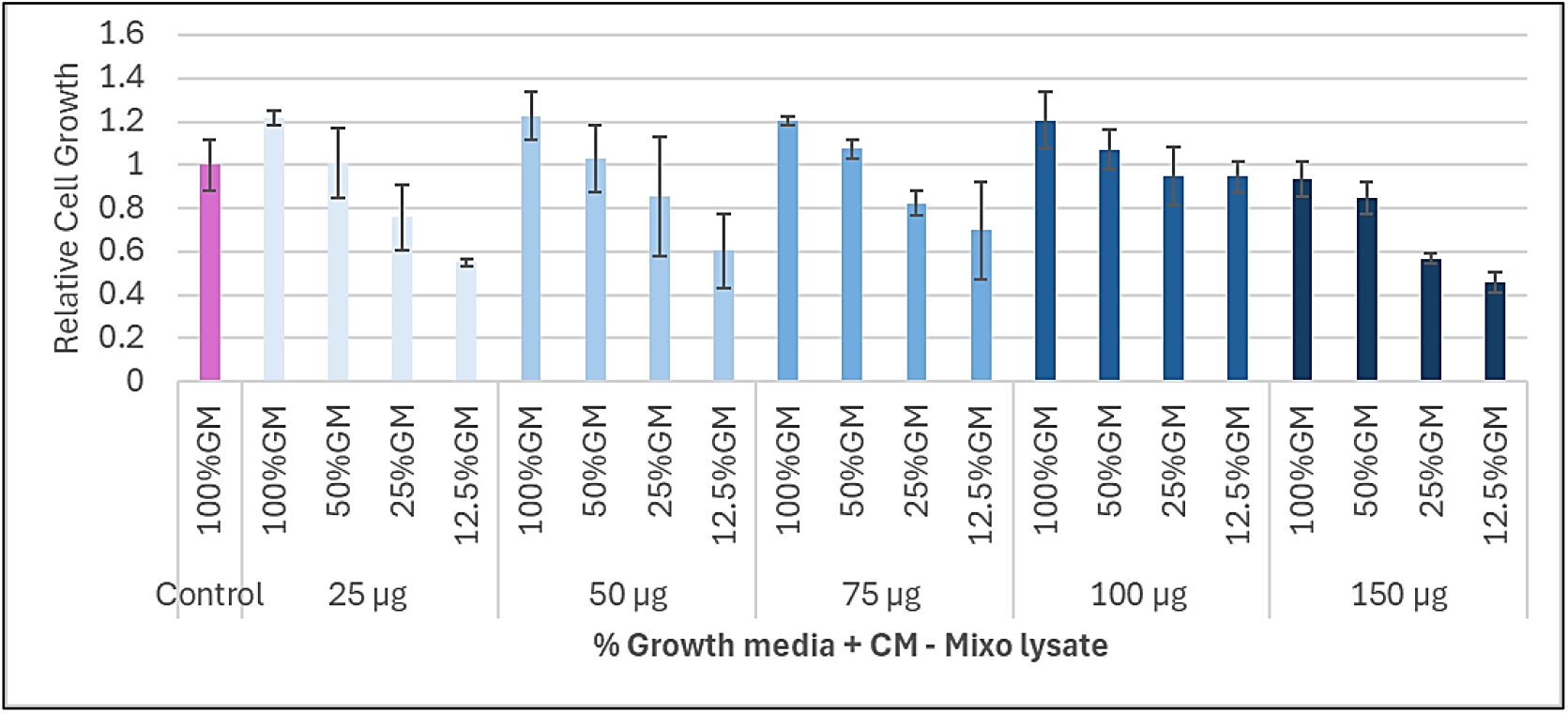
Relative C2C12 cell proliferation in different percentages of growth media supplemented with algae lysate derived from mixotrophic growth in CM spent media (CM-Mixo). Data were derived from up to two independent experiments with three replicates each. Error bars indicate 95% confidence intervals (CI).

**Supplementary Fig. 5:**
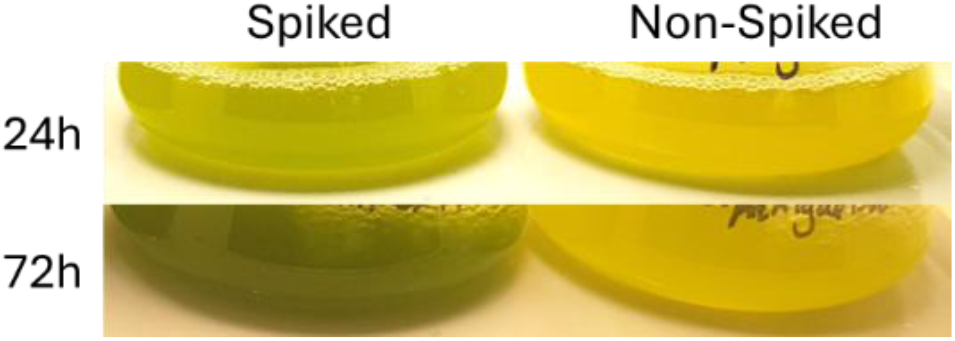
*C*. BDH-1 photoautotrophic growth in twice recycled spent CM media spiked with 10% inorganic algal media (left) and non-spiked (right).

**Supplementary Fig. 6:**
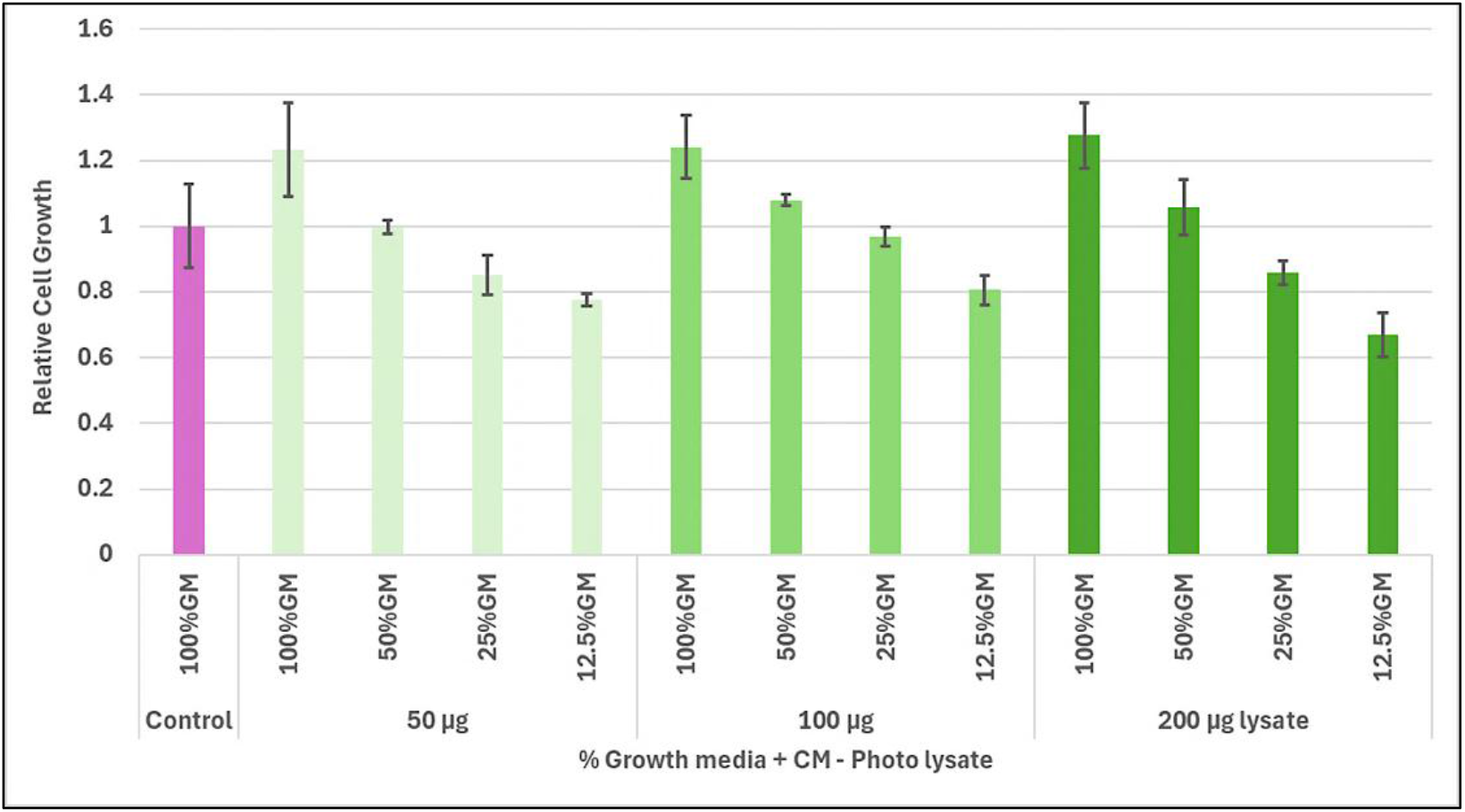
Relative C2C12 cell proliferation in different percentages of growth media supplemented with algae lysate derived from photoautotrophic growth in CM spent media (CM-Photo). Data were derived from up to two independent experiments with three replicates each. Error bars indicate 95% confidence intervals (CI).

